# CHARMER: detecting and harmonizing high-confidence chromatin interactions across tissues and Hi-C protocols

**DOI:** 10.1101/2024.11.25.625258

**Authors:** Simon Cole, Pavel P. Kuksa, Jeffrey Cifello, Otto Valladares, Yuk Yee Leung, Li-San Wang

**Affiliations:** Penn Neurodegeneration Genomics Center, Department of Pathology and Laboratory Medicine, University of Pennsylvania; Embry-Riddle Aeronautical University

## Abstract

**Motivation:** Chromatin conformation capture experiments (CCC), such as Hi-C and Capture Hi-C (CHiC) work to elucidate the three-dimensional organization of the genome and the underlying epigenetic regulatory structures within. CCC experiments produce large amounts of FASTQ sequencing data with a substantial amount of technical noise and require sophisticated computational pipelines in order to extract meaningful results. Large-scale CCC data repositories like 4D Nucleome and ENCODE mostly provide raw contact information but lack annotated, statistically significant interaction data suitable for downstream genetic and genomic analyses.

**Results:** Here, we present CHARMER, an end-to-end pipeline integrated across multiple CCC assay types (HiC, CHiC) which generates statistically significant, harmonized, queryable, chromatin interactions in a consistent BED-like format across cell/tissue types and CCC assays.

**Availability:** CHARMER is freely available at https://bitbucket.org/wanglab-upenn/CHARMER and harmonized chromatin interaction data will be available in the upcoming version of the FILER database (https://lisanwanglab.org/FILER).

## 1 Introduction

Chromatin conformation capture experiments (CCC) work to provide the scientific community with data describing the three-dimensional organization of the genome. Since the advent of the Hi-C protocol in 2009 (Lieberman-Aiden *et al*. 2009), various derivatives of this protocol have been developed which can provide greater enrichment in targeted regions or a more fine-scale resolution with CHiC (Mifsud *et al*. 2015), DNASE Hi-C (Ma *et al*. 2014), or Micro-C (Hsieh *et al*. 2015; Hua *et al*. 2021; Hamley *et al*. 2023). Insights gleaned from these experiments further our understanding of the epigenetic regulatory structures underlying GWAS signals by linking GWAS loci to distal regulatory elements or target genes (Chesi *et al*. 2019; Wachowski *et al*. 2024). CCC experiments generate vast amounts of FASTQ sequencing data and require complex computational analyses to produce biologically meaningful results (Jung *et al*. 2019a), like physically interacting genomic regions, loops, or topologically associated domains (TADs).

Existing large-scale resources in this field such as 4D Nucleome (Reiff *et al*. 2022) and ENCODE (Dunham *et al*. 2012; Luo *et al*. 2020) prioritize the generation, analysis, and distribution of visual Hi-C contact matrices but lack annotated, statistically significant interaction data in consistent formats across other assays. A lack of data harmonization can impede critical downstream analyses like variant-to-gene mapping and cell-type specific target gene inference. Likewise, many datasets don’t include in-depth, standardized metadata and statistics across cell/tissue types and sequencing runs. This can lead to complications in quality control and comparative analysis. Additionally, due to the large amounts of technical noise inherent to CCC experiments (Yardımcı *et al*. 2019), statistical filtering of interactions is critical in providing a high signal-to-noise ratio (Ay, Bailey and Noble 2014; Kaul, Bhattacharyya and Ay 2020). Analyses that include a large number of interactions which likely do not represent true biology could ultimately lead to false conclusions or inaccurate models.

To address these challenges, CHARMER (Chromatin Confirmation Capture Harmonization and Analysis End-to-end fRamework) provides an end-to-end pipeline integrated across multiple chromatin capture assay types (Hi-C, Capture Hi-C) with the goal of generating statistically significant, harmonized, rapidly queryable chromatin interaction data across all the cell/tissue types and biological replicates in a given experiment.

## 2 Implementation or Features

The CHARMER pipeline combines well-established CCC data processing and analysis tools with novel solutions to produce an easy to use fully integrated end-to-end pipeline from raw FASTQ reads to annotated, harmonized, high-confidence interaction sets. CHARMER was primarily developed using Bash/AWK but also utilizes Python3 and R. In terms of features, CHARMER compares favorably to many other CCC pipelines particularly around the data harmonization and annotation features outlined in **Section 2.4 Data Harmonization and Output**. These harmonization, annotation, and indexing features fuel a key use case of CHARMER which is cell/tissue type chromatin interaction survey analyses. A full feature comparison table can be found in **Supplementary Table S1**.

### 2.1 Input

The CHARMER pipeline (**Figure 1**) accepts raw FASTQ sequencing reads across all the cell/tissue types and replicates in a project as input with support for both genome-wide and targeted assays (Hi-C and CHiC). In addition to raw FASTQ reads, the pipeline requires a configuration file as input. This file describes input data characteristics to be used later for metadata generation such as cell and assay types. If CHARMER is running on a CHiC assay, then a BED file marking the captured regions should also be included. A utility script is included to assist in generating these regions using probe sequences or transcription start sites for promoter-capture experiments. After input, CHARMER performs an automatic lift-over step to ensure all data is concordant with the target genomic build (GRCh38/hg38). CHARMER also produces a digested genome file of genomic fragments partitioned by some given restriction enzyme sequence. Various input files required for later steps in the pipeline are also generated at this time.

**Figure 1:**
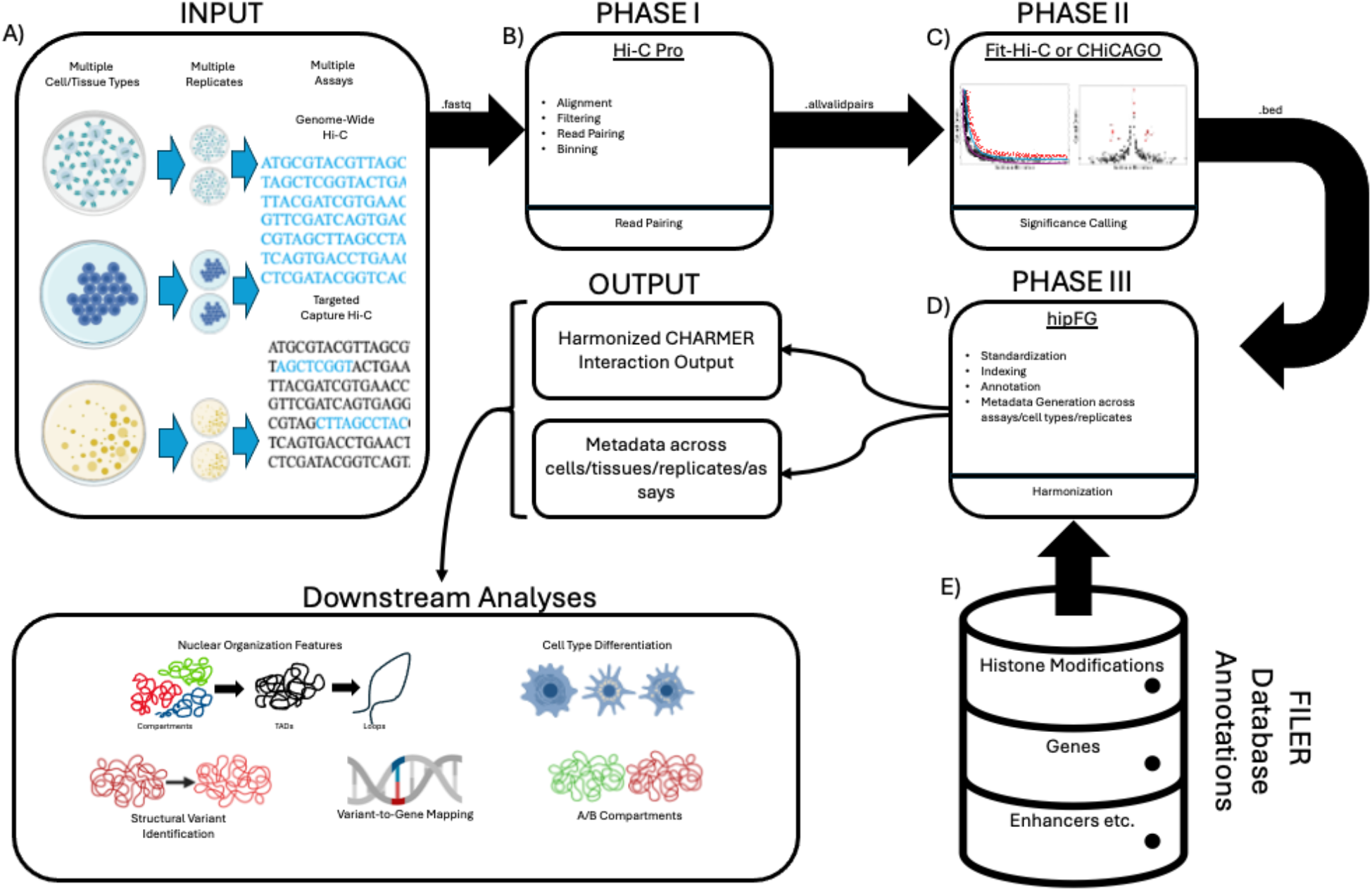
The CHARMER Pipeline Workflow. A) The input to CHARMER. Raw FASTQ sequencing reads are the primary input. CHARMER is designed to accept reads across cell/tissue types with support for multiple CCC assay types (Hi-C and CHiC). B) The read pairing phase of CHARMER. This phase relies heavily upon the Hi-C Pro (Servant *et al*. 2015) pipeline and performs alignment with bowtie2 (Langmead and Salzberg 2012), quality control filtering, pairing, and binning. This phase produces a .allvalidpairs file. C) The significance calling phase of CHARMER. Pairs are assigned statistical significance based on genomic distance and observed contact counts with separate implementations for Hi-C (Ay, Bailey and Noble 2014; Kaul, Bhattacharyya and Ay 2020) and CHiC (Cairns *et al*. 2016). This phase generates a .bed file. D) The data harmonization and annotation phase of CHARMER. Here, data is standardized (Cifello *et al*. 2023) into consistent formatting, GIGGLE indexed (Layer *et al*. 2018), annotated, and robust metadata is generated across replicates and cell/tissue/assay types. E) Annotations for the harmonization phase are retrieved from the FILER (Kuksa *et al*. 2022) functional genomics database. Figure created in part with BioRender.com.

### 2.2 Read Pairing

CHARMER’s read pairing phase is primarily based upon the highly trusted benchmark Hi-C analysis tool, Hi-C pro (Servant *et al*. 2015). The first step in read pairing is aligning the FASTQ sequencing reads to the GRCh38 reference genome, for which CHARMER utilizes bowtie2 (Langmead and Salzberg 2012). CHARMER supports dynamic parallelization by calculating optimal CPU resource allocation and generating SLURM (Yoo, Jette and Grondona 2003) based job submission scripts for the compute heavy alignment steps. After alignment, reads are filtered for quality control (QC) by removing singletons, multi-hit reads, low mapping quality reads, and unmapped reads. The filtered reads are then paired together based on the Hi-C protocol (Belton *et al*. 2012) and filtered further for Hi-C specific QC. Finally, a binning process ensues in order to combine nearby pairs into one interaction between regions where “nearby” is specified as a resolution value. The resultant output from this phase is an allvalidpairs file similar to a BED file in structure.

### 2.3 Significance Calling

The next phase in the CHARMER pipeline is the significance calling of interactions between restriction fragments or bins. CHARMER relies heavily upon two well-established pipelines: Fit-Hi-C for Hi-C (Ay, Bailey and Noble 2014; Kaul, Bhattacharyya and Ay 2020) and CHiCAGO (Cairns *et al*. 2016) for CHiC. The two peak-callers operate on the shared premise of plotting the number of observed interactions between two regions against the genomic distance between them, and from this assigning a statistical significance. Hi-C interactions are called by fitting two non-parametric splines to this plot and removing outliers which lie above the splines to update the model. For CHiC, a background model of expected and observed contact counts following a truncated negative binomial distribution is created. Pairs which then have a contact frequency greater than some expected threshold derived from the model are considered significant. The significance threshold values CHARMER uses are a Fit-Hi-C q-value of < 0.05 or a CHiCAGO score > 5. Interactions are filtered in accordance to these significance thresholds and then saved as BED files.

### 2.4 Data Harmonization and Output

The next phase is data harmonization where significant interactions are processed with the functional genomics (FG) data harmonization pipeline hipFG (Cifello *et al*. 2023), and assigned functional annotations pulled from the FILER (Kuksa *et al*. 2022) FG database. Functional annotations applied to interactions can include gene names, regions, promoters, histone modifications, xQTLs and others. A metadata table with information across cell/tissue/replicate/assay types and data sources is generated. Metadata includes QC results as well as functional trait and interaction statistics like the number of interactions, base-pair coverage, intrachromosomal vs. interchromosomal % etc. Interactions, annotations, and metadata are standardized into consistent formatting, e.g., by normalizing cell type names, assigning broader tissue/cell type categories. Interactions are annotated with source and target gene information, genomic region type (e.g. promoter, intronic, UTR, intergenic). All outputs are also GIGGLE (Layer *et al*. 2018) indexed at this stage to allow for rapid querying. Anchor-based indexing occurs at the cell-type level and a group-based indexing occurs at the experiment level i.e. across cell/tissue/assay types. Tracks can be configured to organize resulting interaction tracks by data source or assay type. CHARMER’s resultant interaction output file is in a BED-like format readily accessible for common downstream analyses like variant-to-gene mapping, survey analyses, or any BEDTOOLS (Quinlan and Hall 2010) operation.

## Results/Discussion

We applied CHARMER to systematically process >400 Chi-C and Hi-C datasets across >27 cell types and tissues (GRCh38/hg38) (Mifsud *et al*. 2015; Jung *et al*. 2019b) (**Supplementary Table S2**). We then used CHARMER’s extensive metadata and harmonized, queryable output to perform multiple analyses. When overlapped with an Alzheimer’s Disease (AD) genome wide association study (Bellenguez *et al*. 2022), processed datasets contained up to 120,000 significant interactions with AD single nucleotide polymorphisms located within them. Additionally, when performing a comparison of the CHiC protocol against the HiC protocol, >300X enrichment of interactions in captured (promoter) regions was observed. In CHiC experiments approximately 92% of significant interactions were between a captured region and a non-captured region, with the remaining 8% being capture-capture. A large number of interchromosomal interactions were observed across cell/tissue and assay types.

The motivations and results outlined in this paper point towards several promising future research directions. First, we plan to expand CHARMER’s compatibility to other impactful CCC assay types like Micro-C (Hsieh *et al*. 2015; Hua *et al*. 2021; Hamley *et al*. 2023), DNase Hi-C (Ma *et al*. 2014) and Single Cell Hi-C (Nagano *et al*. 2013). We also aim to explore binning-free processing approaches similar to HIPPIE (Hwang *et al*. 2015; Kuksa *et al*. 2020) as they can provide fine-scale resolution on interaction site locations through the analysis of read pileups. We also intend to upload more datasets processed with CHARMER to the upcoming version of the FILER FG database.

## Data Availability

CHARMER is freely available at https://bitbucket.org/wanglab-upenn/hicdatapipelines and harmonized chromatin interaction data will be available in the upcoming version of the FILER database (https://lisanwanglab.org/FILER).

## Funding

National Institute on Aging [U24-AG041689, U54-AG052427, U01-AG032984

## Notes

https://bitbucket.org/wanglab-upenn/CHARMER

https://www.niagads.org/FILER2

### Competing Interest Statement

The authors have declared no competing interest.

