## Supplementary Information for "CHARMER: detecting and harmonizing high-confidence chromatin interactions across tissues and Hi-C protocols"

### Supplementary Material

**Supplementary Table S1**: A feature comparison of CHARMER with prominent Hi-C and Capture Hi-C data processing and analysis pipelines

| Feature | | CHARMER This work | HiC-Pro (Servant *et al.* 2015) | Juicer (Servant *et al.* 2015) | Microcket (Zhao *et al.* 2024) | FAN-C (Kruse, Hug and Vaquerizas 2020) | HOMER (Heinz *et al.* 2010) | Hippie (Kuksa *et al.* 2020) | HiC-Bench (Lazaris *et al.* 2017) | HiCUP (Wingett *et al.* 2015) | HiC-Explorer (Wolff *et al.* 2018) |
|---|---|---|---|---|---|---|---|---|---|---|---|
| Raw sequencing data | FASTQ Input | X | X | X | X | X | | X | X | X | X |
| | Alignment | X | X | X | X | X | | X | X | X | X |
| | Read Pairing | X | X | X | X | X | X | X | X | X | X |
| | Dynamic Parallelization | X | | | | X | | | X | X | |
| | Automatic Liftover | X | | | | | | | | | |
| Assays | Genome-Wide Assay Compatibility | X | X | X | X | X | X | X | X | | X |
| | Targeted Assay Compatibility | X | X | | X | | | | | X | X |
| Analysis | Functional Annotation | X | | X | | X | X | X | X | | |
| | Significance-Calling | X | | | | X | X | X | | | X |
| | Standardized Metadata across sources | X | X | | | X | X | | X | X | |
| Harmonization | Indexed & Queryable Output | X | | | | | | | | | |
| | Data Standardization | X | | | | | | | | | |

**Supplementary Table S2**. Chromatin interaction datasets

| Source | Name | Assay Type | # of Cell/Tissue Types | # of Datasets |
|---|---|---|---|---|
| Mifsud et al. 2015 | Mapping long-range promoter contacts in human cells with high-resolution capture Hi-C | Hi-C and CHiC | 2 | 22 |
| Jung et al. 2019b | A Compendium of Promoter-Centered Long-Range Chromatin Interactions in the Human Genome | CHiC | 27 | 383 |
| Reiff et al. 2022 | 4D Nucleome | HiC | 12 | 137 |
